# CIDR: Ultrafast and accurate clustering through imputation for single-cell RNA-Seq data

**DOI:** 10.1101/068775

**Authors:** Peijie Lin, Michael Troup, Joshua W. K. Ho

## Abstract

Most existing dimensionality reduction and clustering packages for single-cell RNA-Seq (scRNA-Seq) data deal with dropouts by heavy modelling and computational machinery. Here we introduce *CIDR* (Clustering through Imputation and Dimensionality Reduction), an ultrafast algorithm which uses a novel yet very simple ‘implicit imputation’ approach to alleviate the impact of dropouts in scRNA-Seq data in a principled manner. Using a range of simulated and real data, we have shown that *CIDR* improves the standard principal component analysis and outperforms the state-of-the-art methods, namely *t-SNE, ZIFA* and *RaceID*, in terms of clustering accuracy. *CIDR* typically completes within seconds for processing a data set of hundreds of cells, and minutes for a data set of thousands of cells. *CIDR* can be downloaded at https://github.org/VCCRI/CIDR.

## Introduction

scRNA-Seq enables researchers to study heterogeneity between individual cells and define cell types from a transcriptomic perspective. One prominent problem in scRNA-Seq data analysis is the prevalence of dropouts, caused by failures in amplification during the reverse-transcription step in the RNA-Seq experiment. The prevalence of dropouts manifests as an excess of zeros and near zero counts in the data set, which has been shown to create difficulties in scRNA-Seq data analysis^1, 2^.

Several packages have recently been developed for the various aspects of scRNA-Seq data analysis, including cell cycle (*cyclone*^3^ and *scLVM*^4^), normalization (*scran*^5^), differential expression analysis (*scde*^2^ and *MAST*^6^) and temporal analysis (*Monocle*^7^), but few perform pre-processing steps such as dimensionality reduction and clustering, which are critical steps for studying cell type heterogeneity.

The state-of-the-art dimensionality reduction package for scRNA-Seq data is *ZIFA*^1^. It implements a modified probabilistic principal component analysis methods that incorporates a zero inflated model to account for dropout events. *ZIFA* uses an iterative expectation-maximization algorithm for inference, which makes it computationally intensive for large scRNA-Seq data sets.

Another package *t-SNE*^8^ is popular among biologists, but it is not designed specifically for scRNA-Seq data and does not address the issue of dropouts. Other recently developed tools such as *BackSPIN*^9^, *pcaReduce*^10^, *SC3*^11^, *SNN-Cliq*^12^, *RaceID*^13^, and *BISCUIT*^14^, were designed to deal optimal clustering of single cells into meaningful groups or hierarchies. Like *ZIFA*, these algorithms usually involve statistical modelling, which themselves require estimation of parameters. These algorithms often make uses of iterative methods to achieve local or global optimal solutions, and hence can be slow when processing large data sets of more than several hundred single cells.

In many practical situations, researchers are interested in fast and intuitive clustering results that they can easily visualise. Principal Component Analysis (PCA) is a common analytical approach for data visualisation for sample heterogeneity, and is often used for dimensionality reduction prior to clustering. Many versions of PCA, such as the implementation *prcomp* in R, is very fast and have been routinely been used for analysing large gene expression data sets. Nonetheless, standard PCA is not designed to take into account of dropouts in scRNA-Seq data. In this work, we aim to develop a fast PCA-like algorithm that takes into account of dropouts.

## Results

**Motivation** We note that PCA is equivalent to performing a Principal Coordinate Analysis (PCoA) on an Euclidean distance matrix derived from the data set. We posit that as long as we can reliably estimate the dissimilarity between every pair of samples (*i.e.,* single cells) in the presence of dropouts, there is no need to explicitly estimate the values of dropouts.

Let’s begin by examining the squared Euclidean distance between the expression profiles of two single cells, 
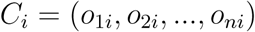
 and 
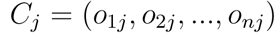
 where *o_ki_* and *o_kj_* represent the gene expression values of gene *k* in cells *C_i_* and *C_j_* respectively:

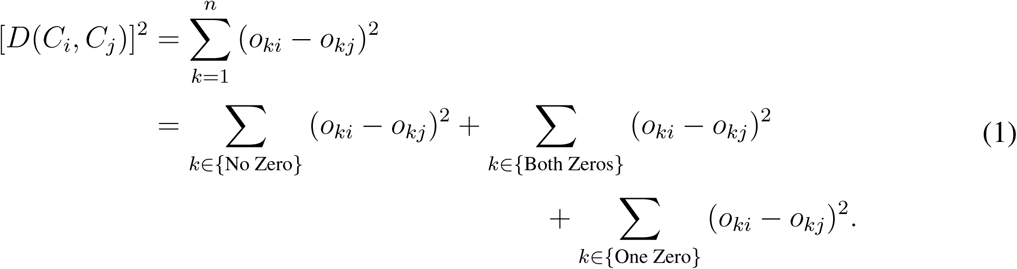

For simplicity, we refer to all zeros in the gene expression data as dropout candidates. In general our argument remains valid even when a dropout candidate is allowed to have near zero values. We note that the squared Euclidean distance in Equation 1 can be arranged as a sum of three sum-of-squares terms. The first term is the sum of squared differences of *o_ki_* and *o_kj_* if they are both non-zero values. This term is not affected by dropout. The second term is the sum of squared differences of *o_ki_* and *o_kj_* if they are both zeros, so this term is zero (or very small if we include near zero values as dropout candidates).

Therefore we observe that the main impact of dropouts comes from the third term, which deals with the case where one value is zero and the other is not. A zero can either represent a lack of gene expression in the ground truth or a dropout event in which a non-zero gene expression value is observed as a zero. If we treat all observed zeros as lack of gene expression (therefore treating the probability of it being a dropout event as zero), which is the case if we directly apply PCA to scRNA-Seq data, this term will tend to be inflated. Nonetheless, it has been observed that the probability of gene expression value being a dropout is inversely correlated with the true expression levels^1,2^. This means a gene with low expression is more likely to become a dropout than a gene with high expression. Using this information, we hypothesise that we can shrink this dropout-induced inflation by imputing the expression value of a dropout candidate in the third term in Equation 1 with its expected value given the dropout probability distribution. This is the motivation behind our new method *CIDR* (Clustering through Imputation and Dimensionality Reduction).

**The *CIDR* algorithm** The *CIDR* algorithm can be divided into the following five steps: (1) Identification of dropout candidates, (2) estimation of the relationship between dropout rate and gene expression levels, (3) calculation of dissimilarity between the imputed gene expression profiles for every pair of single cells, (4) PCoA using the *CIDR* dissimilarity matrix, and (5) clustering using the first few principal coordinates (Supplementary Figure 1).

*CIDR* first performs a logarithmic transformation on the tag per million (TPM). Gene expression for each cell. The distribution of the log-transformed expression values in a scRNA-Seq data set is typically characterised by a strong peak at zero, and one or more smaller non-zero positive peaks representing the expression of expressed genes^6, 15, 16^.

For each cell *C_i_*, *CIDR* finds a sample-dependant threshold *T_i_* that separates the ‘zero peak’ from the rest of the expression distribution; Supplementary Figure 2a shows the distribution of tags for a library in a simulated data set, and the red vertical line indicates the threshold *T_i_*. The entries for cell *C_i_* with an expression of less than *T_i_* are dropout candidates, and the entries with an expression of at least *T_i_* are referred to as ‘expressed’. We call this threshold *T_i_* the ‘dropout candidate threshold’. Note that dropout candidates include true dropouts as well as true low (or no) expressions.

The next step of *CIDR* involves estimating the relationship between dropout probability and gene expression levels. Let *u* be the unobserved true expression of a feature in a cell and let *p(u)* be the probability of it being a dropout. Empirical evidence suggests that *p(u)* is a decreasing function^1, 2^. *CIDR* uses non-linear least squares regression to fit a decreasing logistic function to the data (empirical dropout rate versus average of expressed entries) as an estimate for *p(u)*, illustrated by the ‘Tornado Plot’ Supplementary Figure 2b for the simulated data set. Using the whole data set to estimate *p(u)*, which we denote as 
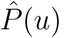
, makes the reasonable assumption that most dropout candidates in the data set are actually dropouts, and allows the sharing of information between genes and cells.

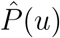
 is used for imputation in the calculation of the *CIDR* dissimilarity matrix. The dropout candidates are treated as missing values and we will now describe *CIDR*’s pairwise ‘implicit’ imputation process. Consider a pair of cells *C_i_* and *C_j_*, and their respective observed expressions *o_ki_* and *o_kj_* for a feature *F_k_*, and let *T_i_* and *T_j_* be dropout candidate thresholds defined as above. Imputation is only applied to dropout candidates, hence the case in which *o_ki_* ≥ *T_i_* and *o_kj_* ≥ *T_j_* requires no imputation. Now consider the case in which one of the two expressions is below *T_i_*, say *o_ki_* < *T_i_* and *o_kj_* ≥ *T_j_*; in this case *o_ki_* needs to be imputed and the imputed value *ô_ki_* is defined as the weighted mean

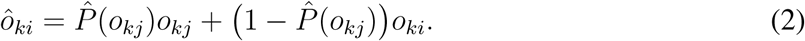

To achieve fast speed in the implementation of the above step, we replace 
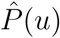
 with a much simpler step function *W(u)*, defined as

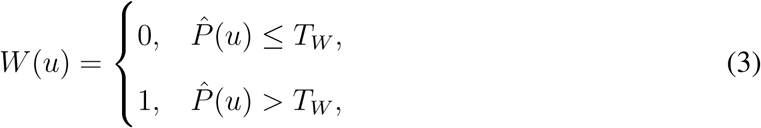

where *T_w_* is by default 0:5. We refer to *W(u)* as the ‘imputation weighting function’ as it gives us the weights in the weighted mean in the imputation, and we refer to the jump of *W(u)*, i.e., 
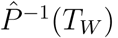
, as the ‘imputation weighting threshold’ (Supplementary Figure 2c). Therefore, the implemented version of Equation (2) is

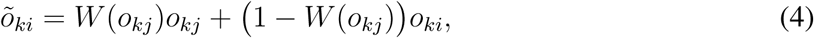

where *õ_ki_* is used as the imputed value of *o_ki_*. Lastly, if *o_ki_* < *T_i_* and *o_kj_* < *T_j_*, we set both *õ_ki_* and *õ_ki_* to be zeros.

We have also implemented *CIDR* directly using 
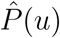
 without the step function simplification. Shown by Tables 1 and 3, the simplification step indeed speeds up the algorithm, and Tables 2 and 3 show that the step does not compromise clustering accuracy.

**Table 1:** Runtime comparison between *CIDR* and four other algorithms. *CIDR* is the default *CIDR* algorithm implementation with step function simplification, while *CIDR* (L) is the implementation with the non-simplified logistic function. The algorithms were run on a standard laptop: 2.8 GHz Intel Core i5 (I5-4308U), 8GB DDR3 RAM). \**RaceID* failed to converge for the mouse brain data set.

| Data set | Size | *CIDR* | *CIDR* (L) | *prcomp* | *t-SNE* | *RaceID* | *ZIFA* |
| --- | --- | --- | --- | --- | --- | --- | --- |
| Pancreatic Islet | 60 | 5.2s | 5.3s | 2.9s | 8.5s | 48.6s | 40.1mins |
| Simulation | 150 | 1.9s | 2.3s | 2.9s | 14.2s | 20.7s | 32.1mins |
| Human Brain | 420 | 6.6s | 8.9s | 13.7s | 1.4mins | 1.5mins | 1.1hours |
| Mouse Brain | 1,800 | 57.9s | 1.1mins | 3.2mins | 23.1mins | 2.5hours* | 1.8hours |

**Table 2:** Clustering accuracy (measured by Adjusted Rand Index) comparison between *CIDR* and four other algorithms. *CIDR* is the default *CIDR* algorithm implementation with step function simplification, while *CIDR* (L) is the implementation with the non-simplified logistic function. *RaceID* failed to converge for the mouse brain data set.

| Data set | Size | *CIDR* | *CIDR* (L) | *prcomp* | *t-SNE* | *RaceID* | *ZIFA* |
| --- | --- | --- | --- | --- | --- | --- | --- |
| Pancreatic Islet | 60 | 0.68 | 0.42 | 0.21 | 0.20 | 0.22 | 0.20 |
| Simulation | 150 | 0.92 | 0.90 | 0.48 | 0.02 | 0 | 0.00 |
| Human Brain | 420 | 0.90 | 0.88 | 0.48 | 0.57 | 0.39 | 0.53 |
| Mouse Brain | 1,800 | 0.52 | 0.37 | 0.26 | 0.62 | 0.37* | 0.32 |

**Table 3:** Runtime and clustering accuracy (measured by Adjusted Rand Index) comparison between *CIDR* and four other algorithms on a simulation data set with 10,000 cells. *CIDR* is the default *CIDR* algorithm implementation with step function simplification, while *CIDR* (L) is the implementation with the non-simplified logistic function. The algorithms except *ZIFA* were run on an AWS ec2 r3.2xlarge instance. \**ZIFA* ran out of memory on the AWS ec2 r3.2xlarge instance, and its runtime was recorded from a run on an AWS ec2 r3.8xlarge instance. \*\**RaceID* didn’t complete after 14 days.

| Simulation (10K) | *CIDR* | *CIDR* (L) | *prcomp* | *t-SNE* | *RaceID* | *ZIFA* |
| --- | --- | --- | --- | --- | --- | --- |
| Time | 44.5mins | 1.5hours | 3.1hours | 21.8hours | >14days | 1.6days* |
| Adjusted Rand Index | 0.99 | 1.00 | 0.99 | 0.00 | N/A** | 0.09 |

Then, the dissimilarity between *C_i_* and *C_j_* is calculated using Equation 1 with the imputed values. We call this imputation approach ‘implicit’, as the imputed value of a particular observed expression of a cell changes each time when it is paired up with a different cell.

Dimensionality reduction is achieved by performing PCoA on the *CIDR* dissimilarity matrix. It is known that clustering performed on the reduced dimensions improves the results^17^. *CIDR* performs hierarchical clustering on the first few principal coordinates, and decides the number of clusters based on the Calinski-Harabasz Index^18^.

**Toy example** Figure 1 shows a toy example that illustrates the effect of dropouts and how *CIDR* can improve clustering in the presence of dropouts. The toy data set consists of eight cells that form two clusters (the red cluster: *c*1-*c*4, and the blue cluster: *c*5-*c*8; Figure 1a). Dropout affects mostly genes with lower expression levels, and hence has a greater impact on cells in the red cluster. Clustering quality can be quantified by the mean squared distance between every pair of cells within a cluster (WC distance) and between clusters (BC distance). The data set is said to have a strong clustering structure if it has low WC distances and high BC distances. In other words, a high ratio of BC/WC distances is an indication of good clustering structure. As illustrated in Fig 1a and 1b, dropout increases both within and between cluster distances. In this case, it also decreases the BC/WC ratio. Using the *CIDR* dissimilarity matrix, we were able to greatly shrink the mean within-cluster distance, while mostly maintain the mean between-cluster distance. In other words, *CIDR* can shrink the within-cluster distances more than the between-cluster distances in a dropout-affected data set. As a result, *CIDR* is able to better preserve the clustering relationship in the original non-dropout data set (Figure 1c).

**Figure 1:**
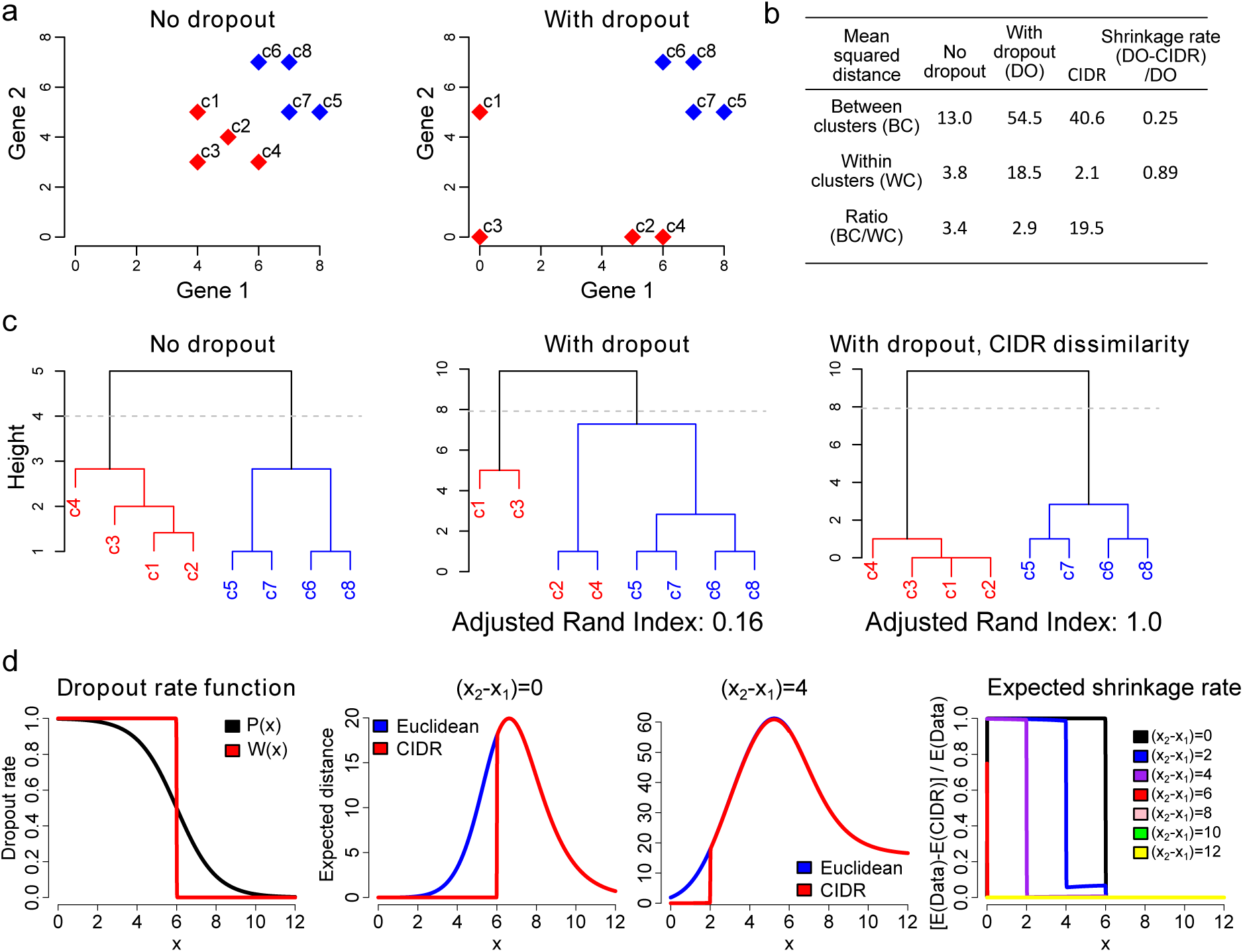
A toy example to illustrate the effect of dropout in scRNA-Seq data on clustering and how *CIDR* can alleviate the effect of dropouts. (a) This toy example consists of eight single cells divided into two clusters (the red cluster and blue cluster. Dropout causes the within-cluster distances among the single cells in the red cluster to increase dramatically, as well as increasing the between cluster distances between single cells in the two clusters. (b) *CIDR* reduces the dropout-induced within-cluster distances while largely maintains the between-cluster distances. (c) The hierarchical clustering results using the original data set (no dropout), the dropout-affected data set, and the dropout-affected data set analysed using *CIDR*.

As a comparison, we have also considered an alternative method in which dropout candidates were imputed to the row mean (IRM) of the expressed entries. This is a straightforward and commonly-used approach for dealing with data with missing values. When applying IRM to our toy data set, we observe that both the between– and within-cluster distances shrink very significantly (Supplementary Figure 3). In fact, in this case IRM shrinks the between-cluster distances a lot more than the within-cluster distances, and therefore dilutes the clustering signal.

This toy example illustrates that the power of *CIDR* comes from its ability to shrink dropout-induced within-cluster distances while largely maintain the between-cluster distances. For theoretical justification, see the **Methods** section.

### Simulation study

For evaluation, we have created a realistic simulated scRNA-Seq data set. We set the number of markers for each cell type low to make it a difficult data set to analyse. Supplementary Figure 2a shows the distribution of tags for one randomly chosen library in this simulated data set. The spike on the left is typical for scRNA-Seq data sets and the tags in this spike are dropout candidates. We have compared *CIDR* with the standard PCA implemented by the R function *prcomp*, two state-of-the-art dimensionality reduction algorithms – *t*-*SNE* and *ZIFA*, and the recently published scRNA-Seq clustering package *RaceID*. As *RaceID* does not perform dimensionality reduction, the first two dimensions output by *t*-*SNE* have been used in the two dimensional visualisation of *RaceID*. Since *prcomp*, *ZIFA* and *t*-*SNE* do not perform clustering, for the purpose of comparison, we have applied the same hierarchical clustering procedure used by *CIDR*. We use the Adjusted Rand Index^19^ to measure the accuracy of clustering.

As shown in Figure 2, the only algorithm that displays three clearly recognisable clusters in the first two dimensions is *CIDR*. *CIDR*’s accuracy in cluster membership assignment is reflected by an Adjusted Rand Index much higher than the other four compared algorithms (Figure 2f). *CIDR* outputs all the principal coordinates as well as a plot showing the proportion of variation explained by each of the principal coordinates (Supplementary Figure 2d).

**Figure 2:**
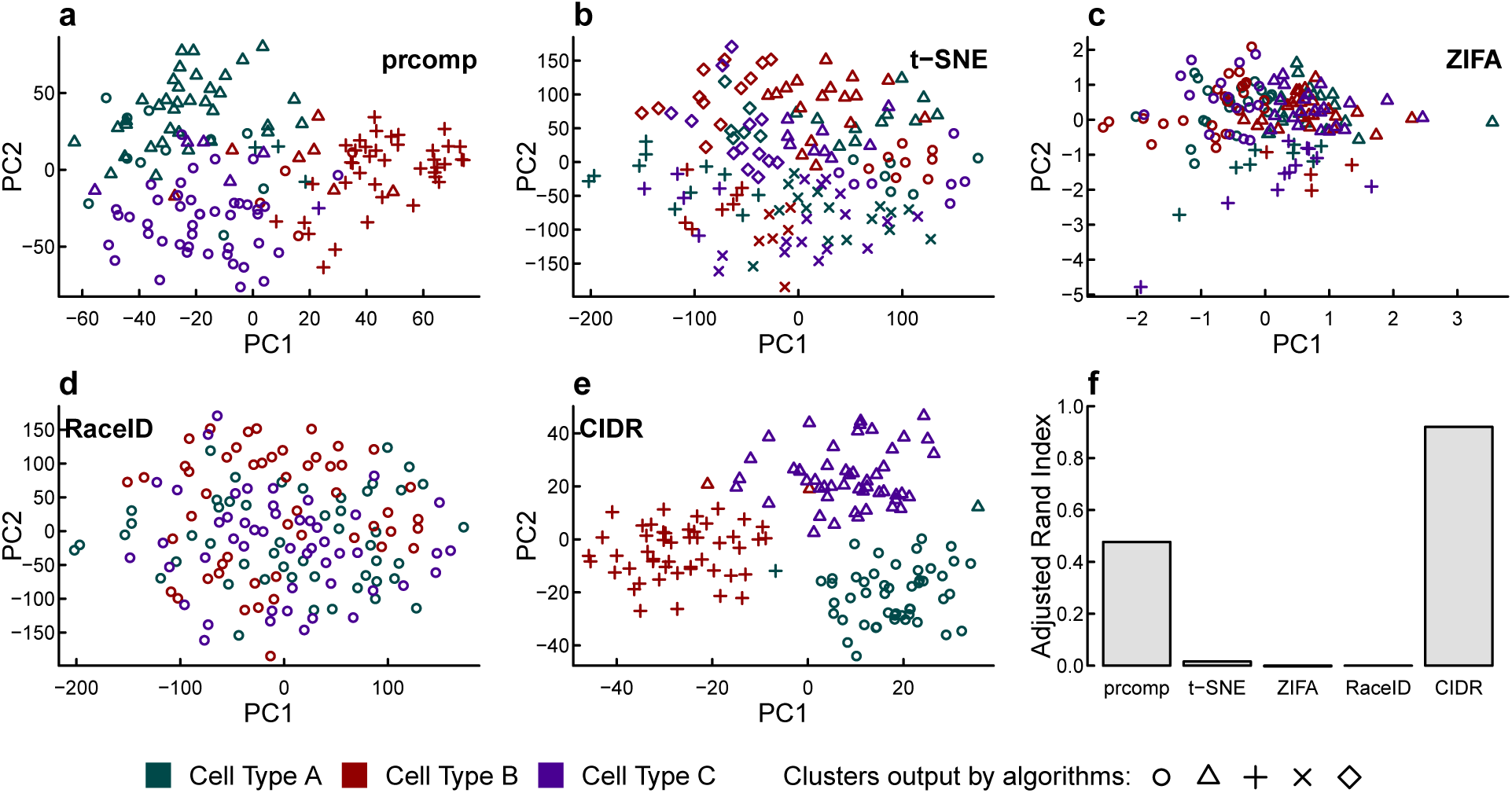
Performance evaluation with simulated data. Simulated scRNA-Seq data set parameters: 3 cell types, 50 cells in each cell type, 20,000 non-differentially expressed features, 150 differentially expressed features and 10 markers for each cell type. The three colors denote the three true cell types; while the different plotting symbols denote the clusters output by each algorithm. (a) ߝ (e) Clustering output by each of the five compared algorithms; (f) Adjusted Rand Index is used to compare the accuracy of the clustering output by each of the compared algorithms.

We perturbed the various parameters in the simulation study to test the robustness of *CIDR* and examine how its performance depends on these parameters. As expected, the Adjusted Rand Index decreases as the dropout level or the number of cell types increases (Supplementary Figures 4a and 4c). However, in cases when the Adjusted Rand Index is low, the performance of *CIDR* can be improved to close to 1 by increasing the number of cells (Supplementary Figures 4b and 4d).

**Scalability of** *CIDR* Given the ever increasing size of scRNA-Seq data sets, and hence the importance of speed of scRNA-Seq data analysis software, we have created a simulated data set of 10,000 cells to test the scalability of *CIDR* and other algorithms. The results are shown in Table 3 - *CIDR* completed the analysis within 45 minutes, which is more than 4-fold faster than the second fastest algorithm *prcomp* (3.1 hours), and many more times faster than *t*-*SNE* (21.8 hours), *ZIFA* (1.6 days), and *RaceID* (did not complete execution within 14 days). In fact, *CIDR* is the only algorithm that completed the analysis within an hour, while achieving a very high clustering accuracy (Adjusted Rand Index = 1).

### Biological data sets

We have applied *CIDR* and the four compared algorithms on three very different biological data sets where the cell types are reported in the original publications. In these studies, cell types were determined through a multi-stage process involving additional information such as cell type molecular signatures. For the purpose of evaluation and comparison, we have applied each of the compared algorithms only once in an unsupervised manner to test how well each algorithm can recover the cell type assignments in the studies.

**Human brain scRNA-Seq data set** Figure 3 shows the comparison results for the human brain scRNA-Seq data set^20^. In this data set there are 420 cells in 8 cell types after we exclude hybrid cells. Determining the number of clusters is known to be a difficult issue in clustering; *CIDR* has managed to identify 7 clusters in the brain data set, which is very close to 8, the number of annotated cell types in this data set. *CIDR* has also identified the members of each cell type largely correctly, as reflected by an Adjusted Rand Index close to 0:9, which is a great improvement over the second best algorithm (Figure 3f). In the two dimensional visualisation by *CIDR* (Figure 3e), the first principal coordinate separates neurons from other cells, while the second principal coordinate separates adult and fetal neurons. Note that *t*-*SNE* is non-deterministic and it outputs dramatically different plots after repeated runs with the same input and the same parameters but with different seed to the random number generator (Supplementary Figure 5).

**Figure 3:**
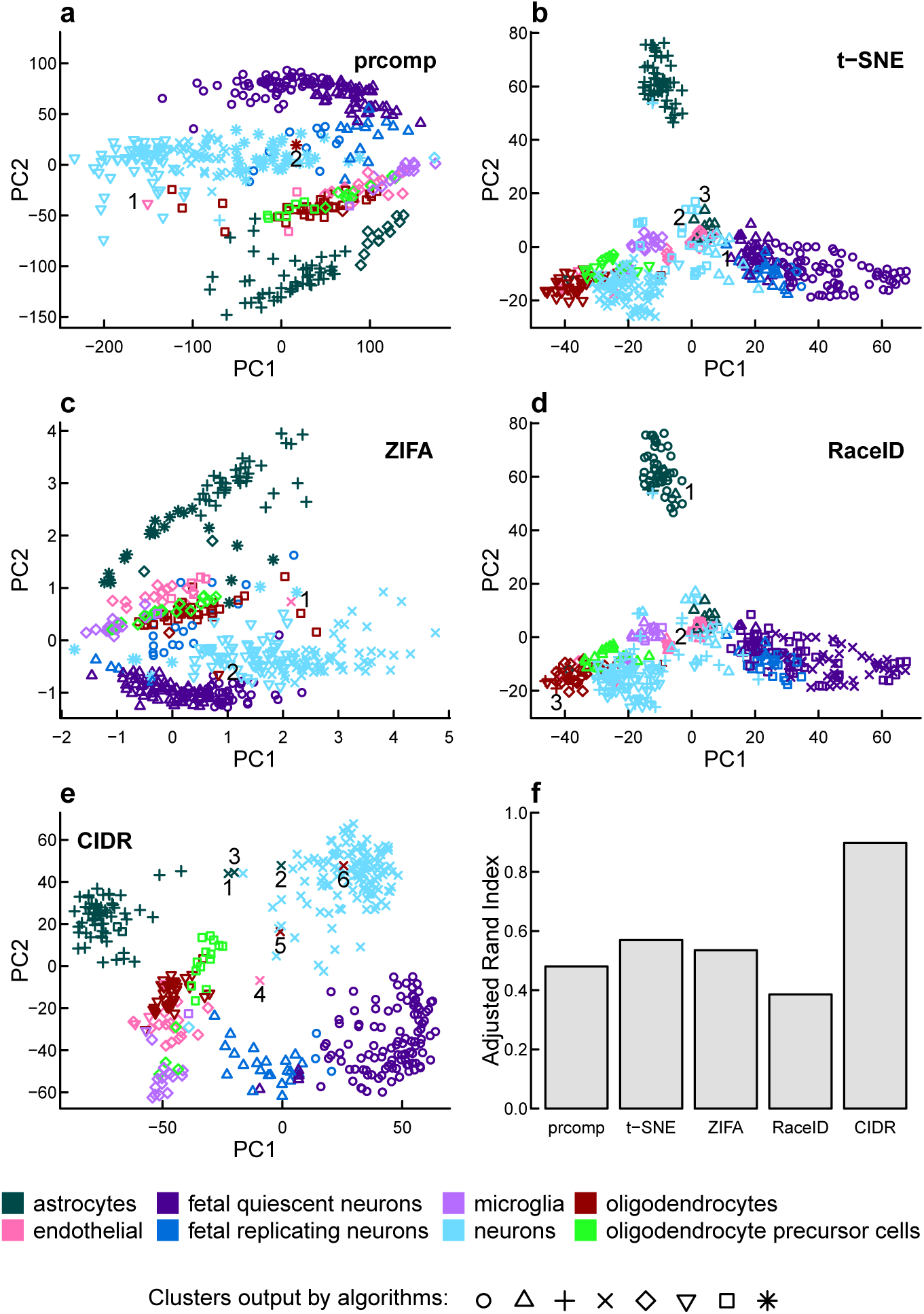
Performance evaluation with the human brain scRNA-Seq data set. In this data set there are 420 cells in 8 cell types after the exclusion of hybrid cells. The different colors denote the cell types annotated by the study^20^; while the different plotting symbols denote the clusters output by each algorithm. (a) – (e) Clustering output by each of the five compared algorithms; (f) Adjusted Rand Index is used to measure the accuracy of the clustering output by each of the compared algorithms. Samples labelled by numbers are disagreements between the annotation and the clustering of the respective algorithm.

*CIDR* allows the user to alter the number of principal coordinates used in clustering and the final number of clusters, specified by the parameters *nPC* and *nCluster* respectively. We altered these parameters and reran *CIDR* on the human brain scRNA-Seq data set to test the robustness of *CIDR* (Supplementary Figure 6). When these parameters are altered from the default values, the clusters output by *CIDR* are still biologically relevant. For instance, 4 is recommended by *CIDR* as the optimal *nPC*, and in the resulting clustering, fetal quiescent neurons and fetal replicating neurons are output as two different clusters (Figure 3e); while when *nPC* is lowered to 2, these two types of cells are grouped as one cluster, i.e., fetal neurons (Supplementary Figure 6a).

We will now use the *CIDR* neuron cluster in the human brain scRNA-Seq data set^20^ as an example to illustrate how to use *CIDR* to discover limitations in the annotation. In Figure 3e the cluster that corresponds best with the annotated neurons is denoted by crosses; there are only six disagreements, marked by 1-6 in Figure 3e, which are denoted by crosses but not annotated as neurons. We use cell type markers from an independent study^21^ to investigate the cause of these disagreements. In Figure 4, these six samples are denoted by ‘*CIDR* 1’, ‘*CIDR* 2’, etc, and as all six samples express neuron markers, *CIDR*’s labels for them are justified. The first five out of these six samples express both neuron markers and the markers of the respective annotated cell types, suggesting that each of these samples contains RNAs from multiple cells, or they are potentially new cell types. The *CIDR* principal coordinates plot (Figure 3e) correctly places these five samples between neurons and the respective annotated cell types. The sixth sample only expresses neuron markers, suggesting a mistake in the annotation, and *CIDR* correctly places this sample in the middle of the neuron cluster. We have carried out the same analysis using *prcomp* and ZIFA, and both methods can only identify ‘*CIDR* 4’ and ‘*CIDR* 6’, marked by 1 and 2 respectively in Figures 3a and 3c. It not not possible to carry out this analysis using *t*-*SNE* or *RaceID*, because they incorrectly group neurons and other cell types in the same clusters. These errors are illustrated in Figures 3b, 3d and 4, in which we can see that cells incorrectly grouped with neurons by *t*-*SNE* and *RaceID*, denoted by ‘t-SNE 1’, ‘t-SNE 2’, etc, have little expression in neuron markers.

**Figure 4:**
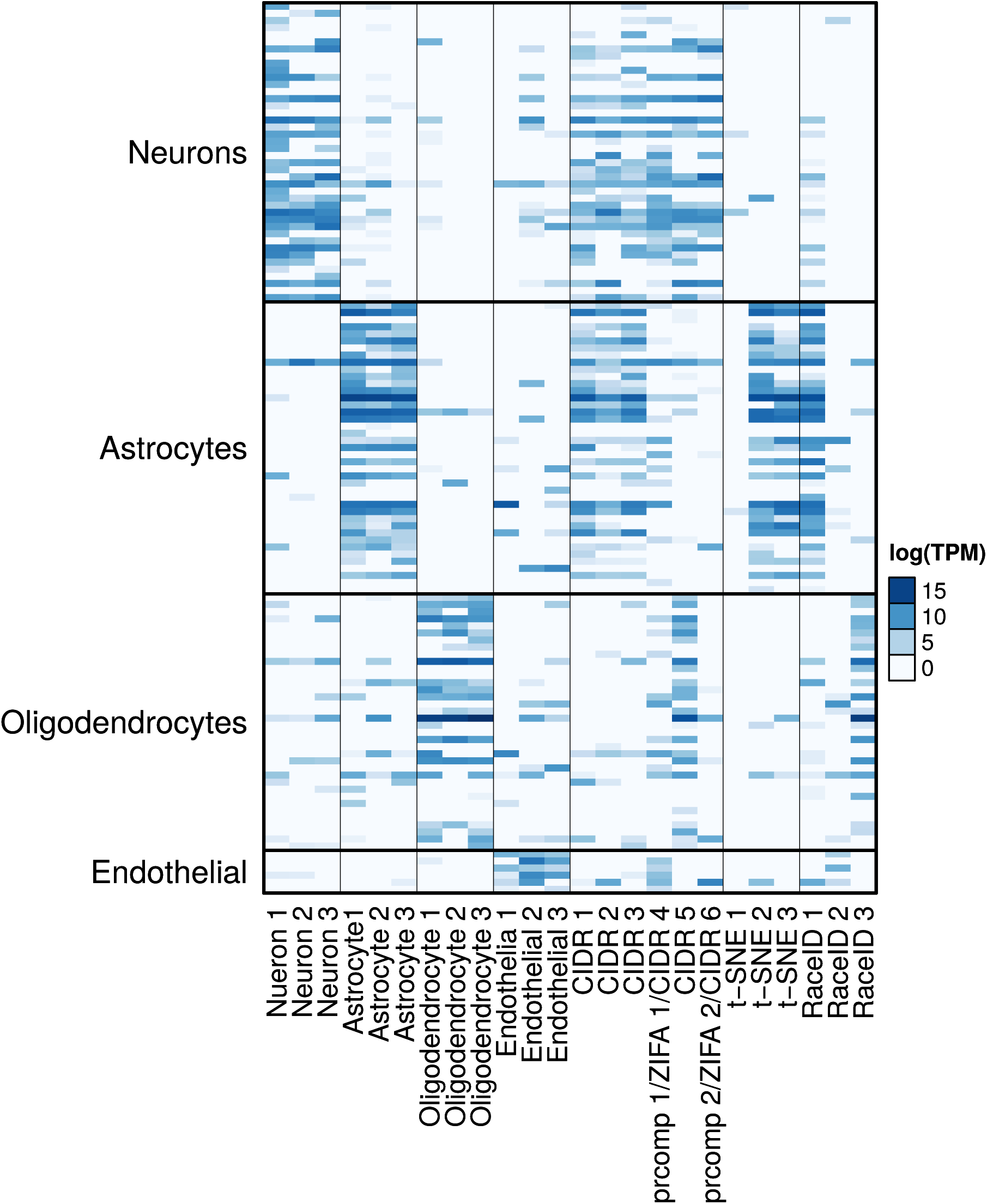
Expression of cell type markers. Four groups of cell type markers from an independent study^21^: neurons, astrocytes, oligodendrocytes and endothelial. The first 12 columns are selected samples which the annotation agrees with the *CIDR* clustering. Columns 13-18 are samples which are not annotated as neurons but clustered with neurons by *CIDR*, *prcomp* or *ZIFA*. Columns 19-24 are selected samples which are not annotated as neurons but clustered with neurons by *t*-*SNE* or *RaceID*.

**Human pancreatic islet scRNA-Seq data set** The human pancreatic islet scRNA-Seq data set^22^ has a smaller number of cells – 60 cells in 6 cell types after we exclude undefined cells and bulk RNA-Seq samples. *CIDR* is the only algorithm that displays clear and correct clusters in the first two dimensions (Figure 5). Regarding clustering accuracy, *CIDR* outperforms the second best algorithm by more than 3-fold in terms of Adjusted Rand Index (Figure 5f).

**Figure 5:**
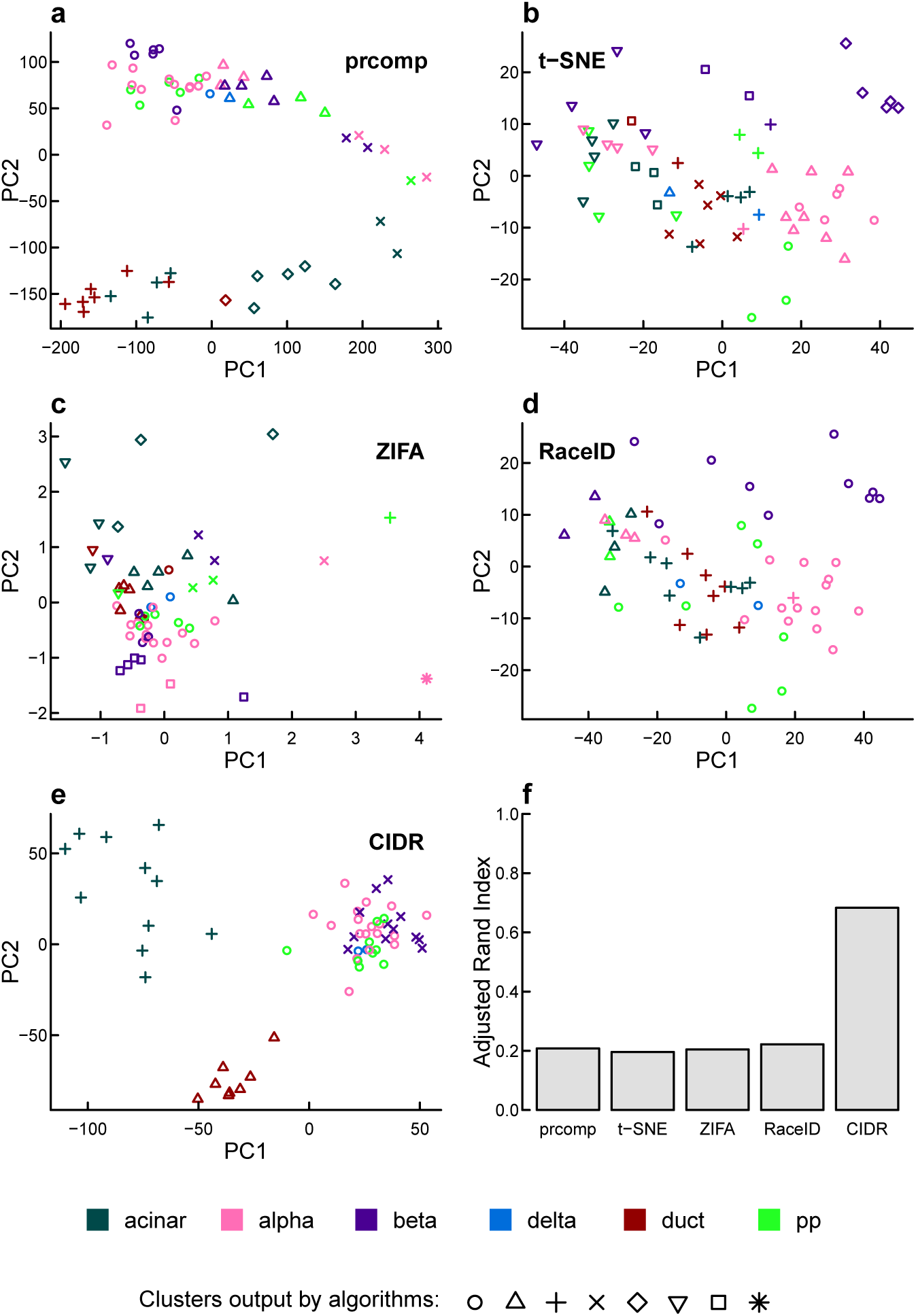
Performance evaluation on the human pancreatic islet scRNA-Seq data set. In this data set there are 60 cells in 6 cell types after the exclusion of undefined cells and bulk RNA-Seq samples. The different colors denote the cell types annotated by the study^22^; while the different plotting symbols denote the clusters output by each algorithm. (a) – (e) Clustering output by each of the five compared algorithms; (f) Adjusted Rand Index is used to measure the accuracy of the clustering output by each of the compared algorithms.

**Mouse brain scRNA-Seq data set** In the mouse brain scRNA-Seq data set^9^ there are 1,800 cells in 7 cell types. Supplementary Figure 7 shows the results of the comparison study using this data set. In this case, *t*-*SNE* achieve the highest Adjusted Rand Index, and this is tightly followed by *CIDR*. Both *t*-*SNE* and *CIDR* perform much better than other tested methods (Table 2 and Supplementary Figure 7), but *CIDR* (1 minute) is significantly faster than *t*-*SNE* (23 minutes) (Table 1). Also, we note that in the original publication^9^ cell-type labels were assigned based on a multi-step procedure involving filtering and applying a modified bi-clustering algorithm, and the clustering results were visualised by *t*-*SNE*.

## Discussion

*CIDR* has ultrafast runtime, which is vital given the rapid growth in the size of scRNA-Seq data sets. The runtime comparison results between *CIDR* and the other four algorithms over five data sets are shown in Tables 1 and 3. On a standard laptop, it takes *CIDR* only seconds to process a data set of hundreds of cells and minutes to process a data set of thousands of cells. It is faster than *prcomp* and all other compared algorithms; in particular, it is more than 50-fold faster than *ZIFA*, which is another dimensionality reduction method that was specifically designed to deal with dropout in scRNA-Seq data analysis.

Data pre-processing steps such as dimensionality reduction and clustering are important in scRNA-Seq data analysis because detecting clusters can greatly benefit subsequent analyses. For example, clusters can be used as covariates in differential expression analysis^6^, or co-expression analysis can be conducted within each of the clusters separately^23^. Certain normalization procedures should be performed within each of the clusters^5^. Therefore, the vast improvement CIDR has over existing tools will be of interest to both users and developers of scRNA-Seq technology.

## Methods

**Dropout candidates** To determine the dropout candidate threshold that separates the first two modes in the distribution of tags (logTPM) of a library, *CIDR* finds the minimum point between the two modes in the density curve of the distribution. The R function *density* is used for kernel density estimation, and the Epanechnikov kernel is used as the smoothing kernel. For robustness, after calculating all the dropout candidate thresholds, the top and bottom 10 percentiles of the thresholds are assigned the 90th percentile and the 10th percentile threshold values respectively. *CIDR* also gives the user the option of calculating the dropout candidate thresholds for only some of the libraries and in this option the median of the calculated thresholds is taken as the dropout candidate threshold for all the libraries.

In the kernel density estimation *CIDR* uses the default bandwidth selection method ‘nrd0’ of the R function density with ‘adjust = 1’. We have varied the ‘adjust’ parameter and re-calculated the Adjusted Rand Indices for both the human brain^20^ and human pancreatic^22^ scRNA-Seq data sets, and Supplementary Figure 8 shows that *CIDR* is robust with respect to this bandwidth adjustment. When the ‘adjust’ parameter is varied from 0:5 to 1:5, the Adjusted Rand Indices for *CIDR* for both the human brain and human pancreatic islet data sets stay much higher than the next best methods; see Figures 3f and Figures 5f.

**Dimensionality reduction** Principal coordinates analysis is performed on the *CIDR* dissimilarity matrix to achieve dimensionality reduction. Because the *CIDR* dissimilarity matrix does not in general satisfy the triangle inequality, the eigenvalues can possibly be negative. This doesn’t matter as only the first few principal coordinates are used in both visualization and clustering, and their corresponding eigenvalues are positive. Negative eigenvalues are discarded in the calculation of the proportion of variation explained by each of the principal coordinates. Some clustering methods require the input dissimilarity matrix to satisfy the triangle inequality. To allow integration with these methods, *CIDR* gives the user the option of Cailliez correction^24^, implemented by the R package *ade4*. The corrected *CIDR* dissimilarity matrix does not have any negative eigenvalues.

***Determining the number of principal coordinates*** *CIDR* implements an algorithm which is a variation of the scree^25^ method to automatically determine the number of principal coordinates used in clustering. *CIDR* outputs a plot that shows the proportion of variation explained by each of the principal coordinates, and the *scree* approach looks for the ‘elbow’ in the curve beyond which the curve flattens.

More specifically, *CIDR* assigns eigenvalues into groups based on the differences in consecutive eigenvalues. A new group is created each time a consecutive difference is greater than a cutoff point determined as a fraction of the largest difference. If the size of the current group exceeds a pre-determined threshold, the sum of sizes of all but the current group is returned as the number of principal coordinates used in clustering.

Users are encouraged to inspect the proportion of variation plot output by *CIDR*, and possibly alter the number of principal coordinates used in clustering.

**Clustering** Hierarchical clustering is performed using the R package *NbClust*. *CIDR*’s default clustering method for hierarchical clustering is ‘ward.D2’^26^, and the number of clusters is decided according to the Calinski-Harabasz Index^18^. The algorithm for cluster number decision is again a variation of the scree^25^ algorithm. More specifically, the algorithm examines the second derivative of the Calinski-Harabasz Index versus the number of clusters curve (Supplementary Figure 2e). Upon user request, *CIDR* can output the Calinski-Harabasz Index versus the number of clusters plot; if needed, the user can overwrite the number of clusters.

**Simulation study** Simulated log tags are generated from a log-normal distribution. For each cell type, an expected library, i.e., the true distribution of log tags, is first generated, and then dropouts and noise are simulated. For each cell type, the expected library includes a small number of differentially expressed features (e.g., genes, transcripts) and markers; by markers we mean features that are expressed in one cell type and zeros in all the other cell types.

A probability function *π(*x*)*, where *x* is an entry in the expected library, is used to simulate dropouts. *π(*x*)* specifies how likely an entry becomes a dropout, so intuitively it should be a decreasing function. In our simulation, we use a decreasing logistic function. The parameters of the logistic function can be altered to adjust the level of dropouts. After the simulation of dropouts, Poisson noise is added to generate the final distribution for each library.

**Biological data** sets Tag tables from three recent scRNA-Seq studies (human brain^20^, human pancreatic islet^22^ and mouse cerebral cortex^9^) were downloaded from the data repository NCBI Gene Expression Omnibus (GSE67835, GSE73727, GSE60361). To ensure good quality, samples with a library size less than 10,000 have been excluded. The raw tag tables were used as the inputs for *CIDR*. For other dimensionality reduction and clustering algorithms, rows with tag sums less than or equal to 10 were deleted. Log tags, with base 2 and prior count 1, were used as the inputs for *ZIFA*, as suggested by the *ZIFA* documentation. Data sets transformed by logTPM were used as inputs for *prcomp* and *t*-*SNE*.

**Theoretical justification** Here we show that *CIDR* always shrink the expect distance between two dropout-affected samples (*i.e.,* single cells), and has a higher expected shrinkage rate for within-cluster distances than between cluster distances. It is this property that ensures that *CIDR* dissimilarity matrix better preserves the clustering structure in the data set.

For simplicity of discussion, let’s assume that dropouts are zeros. We will now explain why imputation by Equation (2) in the main text improves clustering.

Suppose that a particular feature *F* has true expression level *x*_1_, *x*_2_, and *x*_3_ for three cells *C*_1_, *C*_2_, and *C*_3_ respectively. Let’s assume *x*_1_ ≤ *x*_2_ ≤ *x*_3_. Let *P* be the true dropout probability function, and 
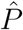
 be the empirically estimated dropout probability function used in *CIDR*. Both 
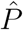
. and P are monotonically decreasing functions, and satisfy 
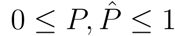

The true dissimilarity between *C*_1_ and *C*_2_ contributed by feature *F* is

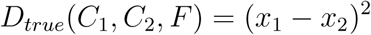

In the presence of dropouts in the observed data, the expected value of dissimilarity between *C_1_* and *C_2_* contributed by feature *F* is

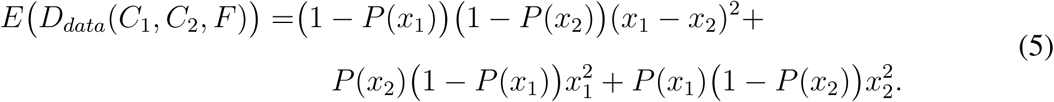

Meanwhile the expected value of the *CIDR* dissimilarity between *C_1_* and *C_2_* contributed by feature *F* is

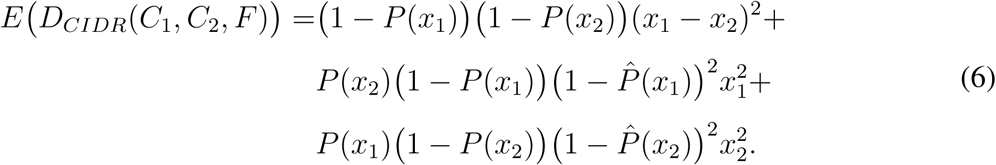

Comparing Equation (5) and Equation (6), it is clear that the only difference is the presence of the factor 
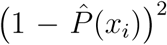
 in the last two terms. Since 
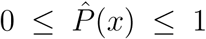
, we can deduce that 
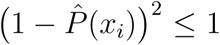
, which means 
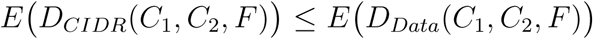
 for pair of cells *C_1_* and *C_2_*. This demonstrates that *CIDR* shrinks the expected distance between two points in the presence of dropouts.

Furthermore, let’s consider the expected rate of shrinkage between *C_1_* and *C_2_* contributed by feature *F*,

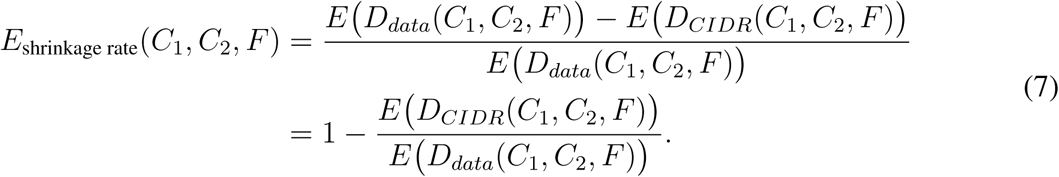

Let’s consider 
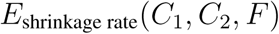
 and 
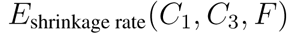
 Since *CIDR* always shrink the expected distance between two points, and that 
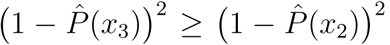
 our intuition is that 
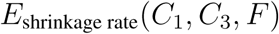
 is likely smaller than or equal to 
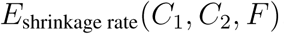
. In other words, we hypothesise that the shrinkage rate between two closer points is larger than or equal to the shrinkage rate between two points that are further apart. To prove this property is algebraically very complex, so we have conducted an extensive computational study on the rate of shrinkage; Supplementary Figure 9 shows that for a variety of monotonically decreasing *P* and 
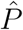
 and for any fixed *x_1_*, the expected rate of shrinkage becomes smaller when *x_2_* becomes larger. In particular, Supplementary Figure 9f shows the case when 
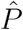
 is a step function. We observe that in all tested cases our hypothesis holds. Therefore we are satisfied that in practice *CIDR* shrinks within-cluster distances more than between-cluster distances due to this differential shrinkage rate property.

## Competing Interests

The authors declare that they have no competing interests.

## Funding

This work is supported in part by the New South Wales Ministry of Health, the Human Frontier Science Program (RGY0084/2014), the National Health and Medical Research Council of Australia (1105271), the National Heart Foundation of Australia, and the Amazon Web Services (AWS) Cloud Credits for Research.

## Acknowledgements

Willem Van Der Byl contributed to parts of the code in the *CIDR* package.

## Additional Files

**Additional file 1 – Supplementary Figures** Supplementary Figures 1 – 9.

## References

1. Pierson, E. & Yau, C. ZIFA: Dimensionality reduction for zero-inflated single-cell gene expression analysis. Genome Biology 16, 1–10 (2015).

2. Kharchenko, P. V., Silberstein, L. & Scadden, D. T. Bayesian approach to single-cell differential expression analysis. Nature Methods 11, 740–742 (2014).

3. Scialdone, A. et al. Computational assignment of cell-cycle stage from single-cell transcriptome data. Methods 85, 54–61 (2015).

4. Buettner, F. et al. Computational analysis of cell-to-cell heterogeneity in single-cell RNA-sequencing data reveals hidden subpopulations of cells. Nature Biotechnology 33, 155–160 (2015).

5. Lun, A. T., Bach, K. & Marioni, J. C. Pooling across cells to normalize single-cell RNA sequencing data with many zero counts. Genome Biology 17, 1 (2016).

6. Finak, G. et al. MAST: a flexible statistical framework for assessing transcriptional changes and characterizing heterogeneity in single-cell RNA sequencing data. Genome Biology 16, 1–13 (2015).

7. Trapnell, C. et al. The dynamics and regulators of cell fate decisions are revealed by pseudotemporal ordering of single cells. Nature Biotechnology 32, 381–386 (2014).

8. Maaten, L. & Hinton, G. Visualizing data using t-SNE. Journal of Machine Learning Research 9, 2579–2605 (2008).

9. Zeisel, A. et al. Cell types in the mouse cortex and hippocampus revealed by single-cell RNA-seq. Science 347, 1138–1142 (2015).

10. Zurauskiene, J. & Yau, C. pcaReduce: hierarchical clustering of single cell transcriptional profiles. BMC Bioinformatics 17, 140 (2016).

11. Kiselev, V. Y. et al. SC3-consensus clustering of single-cell RNA-Seq data. bioRxiv 036558 (2016).

12. Xu, C. & Su, Z. Identification of cell types from single-cell transcriptomes using a novel clustering method. Bioinformatics 31, 1974–80 (2015).

13. Grün, D., et al. Single-cell messenger RNA sequencing reveals rare intestinal cell types. Nature 525, 251–255 (2015).

14. Prabhakaran, S., Azizi, E. & Pe’er, D. Dirichlet process mixture model for correcting technical variation in single-cell gene expression data. In Proceedings of The 33rd International Conference on Machine Learning, 1070–1079 (2016).

15. McDavid, A. et al. Modeling bi-modality improves characterization of cell cycle on gene expression in single cells. PLoS Computational Biology 10, e1003696 (2014).

16. Bacher, R. & Kendziorski, C. Design and computational analysis of single-cell RNA-sequencing experiments. Genome Biology 17, 1 (2016).

17. Ronan, T., Qi, Z. & Naegle, K. M. Avoiding common pitfalls when clustering biological data. Science Signaling 9, re6 (2016).

18. Caliński, T. & Harabasz, J. A dendrite method for cluster analysis. Communications in Statistics 3, 1–27 (1974).

19. Hubert, L. & Arabie, P. Comparing partitions. Journal of Classification 2, 193–218 (1985).

20. Darmanis, S. et al. A survey of human brain transcriptome diversity at the single cell level. Proceedings of the National Academy of Sciences 112, 7285–7290 (2015).

21. Cahoy, J. D. et al. A transcriptome database for astrocytes, neurons, and oligodendrocytes: a new resource for understanding brain development and function. The Journal of Neuroscience 28, 264–278 (2008).

22. Li, J. et al. Single-cell transcriptomes reveal characteristic features of human pancreatic islet cell types. EMBO Reports 17, 178–187 (2016).

23. Trapnell, C. Defining cell types and states with single-cell genomics. Genome Research 25, 1491–1498 (2015).

24. Cailliez, F. The analytical solution of the additive constant problem. Psychometrika 48, 305–308 (1983).

25. Cattell, R. B. The scree test for the number of factors. Multivariate behavioral research 1, 245–276 (1966).

26. Murtagh, F. & Legendre, P. Ward’s hierarchical agglomerative clustering method: Which algorithms implement Ward’s criterion? Journal of Classification 31, 274–295 (2014).

